# Massive crossover elevation via combination of *HEI10* and *recq4a recq4b* during Arabidopsis meiosis

**DOI:** 10.1101/159764

**Authors:** Heïdi Serra, Christophe Lambing, Catherine H. Griffin, Stephanie D. Topp, Mathilde Séguéla-Arnaud, Joiselle Fernandes, Raphaël Mercier, Ian R. Henderson

## Abstract

During meiosis homologous chromosomes undergo reciprocal crossovers, which generate genetic diversity and underpin classical crop improvement. Meiotic recombination initiates from DNA double strand breaks, which are processed into single-stranded DNA that can invade a homologous chromosome. The resulting joint molecules can ultimately be resolved as crossovers. In Arabidopsis, competing pathways balance the repair of ∼100–200 meiotic DSBs into ∼10 crossovers per meiosis, with the excess DSBs repaired as non-crossovers. In order to bias DSB repair towards crossovers, we simultaneously increased dosage of the pro-crossover E3 ligase gene *HEI10* and introduced mutations in the anti-crossover helicase genes *RECQ4A* and *RECQ4B*. As *HEI10* and *recq4a recq4b* increase interfering and non-interfering crossover pathways respectively, they combine additively to yield a massive meiotic recombination increase. Interestingly, we also show that increased *HEI10* dosage increases crossover coincidence, which indicates an effect of *HEI10* on interference. We also show that patterns of interhomolog polymorphism and heterochromatin drive recombination increases towards the sub-telomeres in both *HEI10* and *recq4a recq4b* backgrounds, while the centromeres remain crossover-suppressed. These results provide a genetic framework for engineering meiotic recombination landscapes in plant genomes.

## Main text

Meiosis is a conserved cell division required for eukaryotic sexual reproduction, during which a single round of DNA replication is coupled to two rounds of chromosome segregation, generating haploid gametes^1,2^. Homologous chromosomes pair and recombine during prophase of the first meiotic division, which can result in reciprocal exchange, termed crossover^1,2^. Crossovers have a major effect on sequence variation in populations and create genetic diversity. Meiotic recombination is also an important tool used during crop breeding to combine useful variation. However, crossover numbers are typically low, ∼1–2 per chromosome per meiosis, and can show restricted chromosomal distributions, which limit crop improvement. For example, recombination is suppressed in large regions surrounding the centromeres of many crop species^3^. In this work we sought to use our understanding of meiotic recombination pathways to genetically engineer super-recombining Arabidopsis.

Meiotic recombination initiates from DNA double strand breaks (DSBs), induced by SPO11 transesterases, which act in topoisomerase-VI-like complexes^1,4^ (Fig. 1a). During catalysis SPO11 becomes covalently bound to target site DNA and is liberated by endonucleolytic cleavage by the MRN (MRE11-RAD50-NBS1) complex and COM1^4^. Simultaneously, exonucleases generate 3ʹ-overhanging single stranded DNA (ssDNA), approximately 100-1000s of nucleotides in length^4^ (Fig. 1a). Resected ssDNA is bound by RAD51 and DMC1 RecA-like proteins, which promote invasion of a homologous chromosome and the formation of a displacement loop (D-loop)^5^ (Fig. 1a). Stabilization of the D-loop can occur by template-driven DNA synthesis from the invading 3’-end^4,6^ (Fig. 1a). Strand invasion intermediates may then progress to second-end capture and formation of a double Holliday junction (dHJ), which can be resolved as a crossover or non-crossover, or undergo dissolution^1,4,6^ (Fig. 1a).

**Figure 1.**
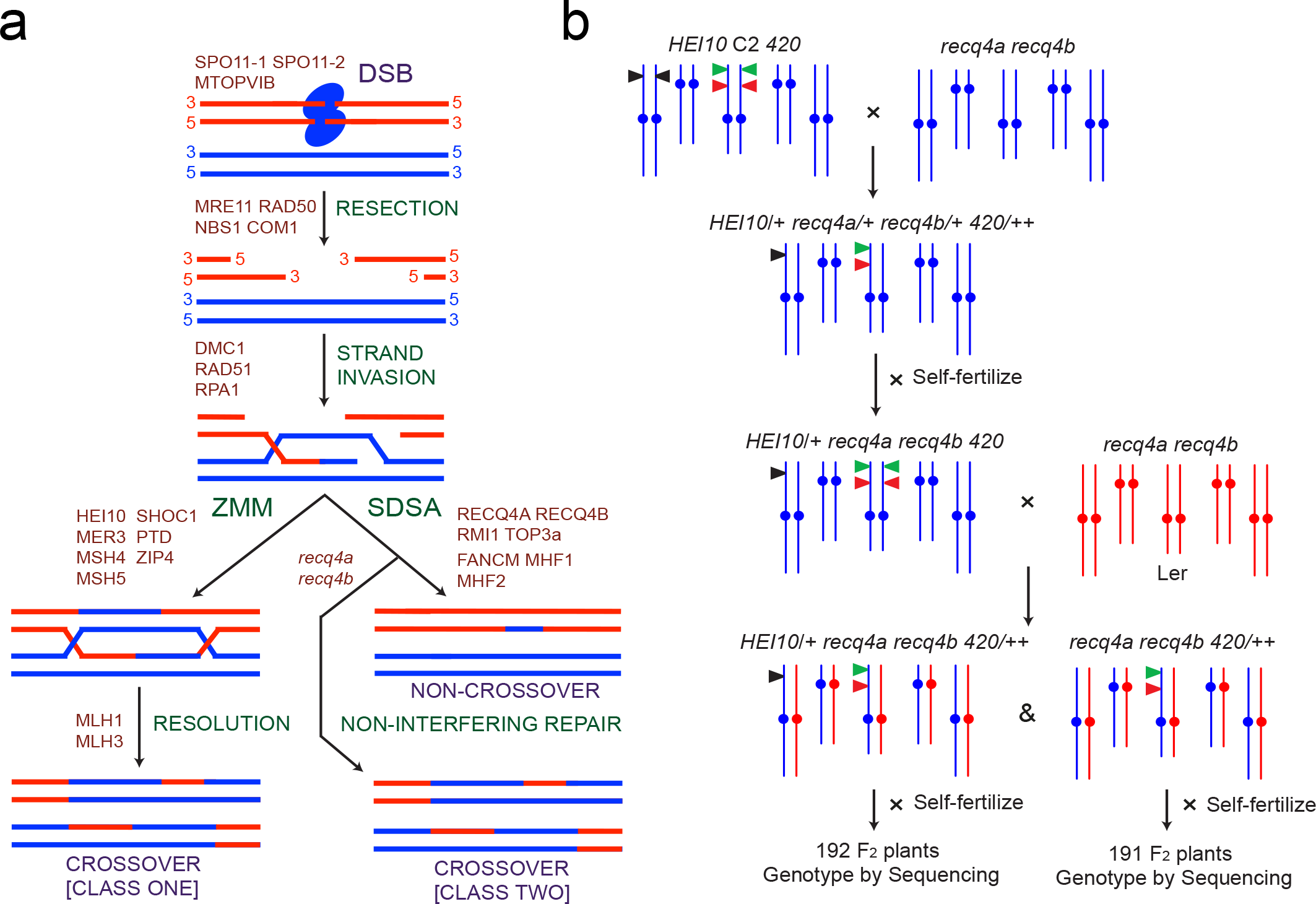
Combining increased *HEI10* dosage with *recq4a recq4b* mutations in order to elevate meiotic crossovers. **a.** Schematic diagram showing a subset of pathways acting during Arabidopsis meiotic recombination. Homologous chromosomes are indicated as red and blue DNA duplexes. The remaining two sister chromatid duplexes are not shown for simplicity. Recombination pathways (green) and factors acting within them (red) are printed alongside chromosomes. **b.** Crossing diagram showing the generation of Col/Ler F_1_ plants that were *recq4a recq4b* mutant, with and without *HEI10* (black triangles). F_2_ progeny from these plants were analysed by genotyping-by-sequencing to map crossover locations. These populations were compared to wild type F_2_ progeny and to a previously reported *HEI10* F_2_ population^12^.

The conserved ZMM pathway acts to promote formation of the majority of crossovers in plants, which are known as class I^1,6,7^ (Fig. 1a). Mutations in ZMM genes severely reduce Arabidopsis crossover frequency, causing univalent chromosome segregation at metaphase I, aneuploid gametes and infertility^1,8,9^. Importantly, ZMM-dependent crossovers show interference, where double crossover events are spaced out more widely than expected by chance^10,11^. The ZMM pathway in plants includes the MSH4/MSH5 MutS-related heterodimer, MER3 DNA helicase, SHORTAGE OF CROSSOVERS1 (SHOC1) XPF nuclease, PARTING DANCERS (PTD), ZIP4/SPO22, HEI10 E3 ligase and the MLH1/MLH3 MutL-related heterodimer^1,6,7^ (Fig. 1A). Within the ZMM pathway the *HEI10* E3 ligase gene shows dosage sensitivity, with additional copies being sufficient to increase crossovers throughout euchromatin^12^. A minority of crossovers in plants, known as class II, do not show interference and are formed by a different MUS81-dependent pathway^13,14^.

From cytological measurement of Arabidopsis DSB-associated foci (e.g. *γ*H2A.X, RAD51 and DMC1) along meiotic chromosomes, it is estimated that between 100–200 breaks initiate per nucleus^15–17^. However, only ∼10 crossovers typically form throughout the genome^18–21^, indicating that anti-crossover pathways prevent maturation of the majority of initiation events into crossovers. Indeed, genetic analysis has identified at least three distinct anti-crossover pathways in Arabidopsis: (i) the FANCM DNA helicase and MHF1 and MHF2 co-factors^22–24^, (ii) the AAA-ATPase FIDGETIN-LIKE1^25^ and (iii) the RTR complex of RECQ4A, RECQ4B DNA helicases, TOPOISOMERASE3α and RMI1^26–28^ (Fig. 1a). For example, *recq4a recq4b* mutants show highly increased non-interfering crossovers when assayed in specific intervals^27^ (Fig. 1a). This is thought primarily to result from a failure to dissolve interhomolog strand invasion events, which are alternatively repaired by the non-interfering crossover pathway(s)^22,25,27^. As combining mutations between these pathways, for example *fancm fidgl1*, leads to additive crossover increases they reflect parallel mechanisms^25^. Hence, during meiosis competing pathways act on SPO11-dependent DSBs to balance crossover and non-crossover repair outcomes (Fig. 1a).

In this work we explore the functional relationship between ZMM pro-crossover and RECQ4 anti-crossover meiotic recombination pathways. Using a combination of increased *HEI10* dosage and *recq4a recq4b* mutations, we observe a massive, additive increase in crossover frequency throughout the chromosome arms. Surprisingly, we observe that increased *HEI10* dosage causes increased crossover coincidence, indicating an effect on interference. We show that *HEI10* and *recq4a recq4b* crossover increases are biased towards regions of low interhomolog divergence, distal from centromeric heterochromatin. Hence, both genetic and epigenetic information likely constrain the activity of meiotic recombination pathways.

### Combination of *HEI10* and *recq4a recq4b* massively elevates crossover frequency

Crossover increases in *HEI10* and *recq4a recq4b* represent mechanistically distinct effects via class I and class II crossover repair pathways (Fig. 1a). We therefore sought to test whether combining these genetic backgrounds would cause further increases in crossover frequency. We previously showed that transgenic line ‘*C2*’ carrying additional *HEI10* copies, shows a ∼2-fold increase in crossovers genome-wide, compared with wild type^12^ (Table 1). We therefore crossed *HEI10* line *C2* to *recq4a recq4b* double mutants, in the Col genetic background^12,27,29^ (Fig. 1b). A previous genetic screen isolated an EMS allele of *recq4a* in Ler^27^. As Ler carries a natural premature stop codon in *recq4b*^27^, this provides a *recq4a recq4b* double mutant in Ler (Fig. 1b). These lines were crossed and F_1_ progeny identified that were heterozygous for Col/Ler polymorphisms, *recq4a recq4b* mutant and with or without *HEI10* (Fig. 1b). These F_1_ plants were then used to generate Col/Ler F_2_ for crossover analysis (Fig. 1b).

**Table 1.** Crossovers in wild type, *HEI10, recq4a recq4b* and *HEI10 recq4a recq4b* Col/Ler F_2_ populations. Crossover numbers identified by genotyping-by-sequencing in the indicated F2 populations are listed. The number of individuals analysed per population is indicated in the *n* row. Absolute crossover (COs) numbers are reported, as well as mean crossovers per F_2_ individual, by chromosome and in total. The CO StDev row shows the standard deviation in total crossover numbers per F2 for each genotype.

|  | GENOTYPE |  |  |  |  |  |  |  |
| --- | --- | --- | --- | --- | --- | --- | --- | --- |
|  | Col/Ler |  | *HEI10*<br>Col/Ler |  | *recq4a recq4b*<br>Col/Ler |  | *HEI10*<br>*recq4a recq4b*<br>Col/Ler |  |
| Chromosome | COs | COs/F<sub>2</sub> | COs | COs/F<sub>2</sub> | COs | COs/F<sub>2</sub> | COs | COs/F<sub>2</sub> |
| 1 | 437 | 1.8 | 765 | 4.0 | 1259 | 6.6 | 1641 | 8.5 |
| 2 | 326 | 1.3 | 525 | 2.7 | 799 | 4.2 | 1001 | 5.2 |
| 3 | 345 | 1.4 | 546 | 2.8 | 993 | 5.2 | 1195 | 6.2 |
| 4 | 309 | 1.3 | 417 | 2.2 | 576 | 3.0 | 663 | 3.5 |
| 5 | 423 | 1.7 | 675 | 3.5 | 1156 | 6.1 | 1448 | 7.5 |
| Total | 1840 | 7.5 | 2928 | 15.3 | 4783 | 25.0 | 5948 | 31.0 |
| *n* | 245 |  | 192 |  | 191 |  | 192 |  |
| CO StDev | 2.26 |  | 3.41 |  | 6.52 |  | 6.59 |  |

During crossing we maintained the *420* FTL crossover reporter within our lines, which allows measurement of genetic distance in a ∼5.1 Mb sub-telomeric region on chromosome 3^30,31^ (Figs. 1b, 2a and Supplementary Table 1). This showed that *HEI10, recq4a recq4b* and *HEI10 recq4a recq4b* all significantly increase *420* crossover frequency in Col/Ler backgrounds, by 2.7, 3.3 and 3.7-fold, respectively (*X^2^ P*=2.73×10^−175^, *P*=4.92×10^−212^ and *P*=2.80×10^−226^) (Fig. 2a and Supplementary Table 1). However, it is notable that *420* genetic distance reached 47 cM in *HEI10 recq4a recq4b*, which is close to the maximum observable recombination frequency for linked markers (i.e. 50 cM) (Fig. 2a and Supplementary Table 1). We next used genotyping-by-sequencing (GBS) to generate genome-wide, high-resolution maps of crossover distributions in these backgrounds.

**Figure 2.**
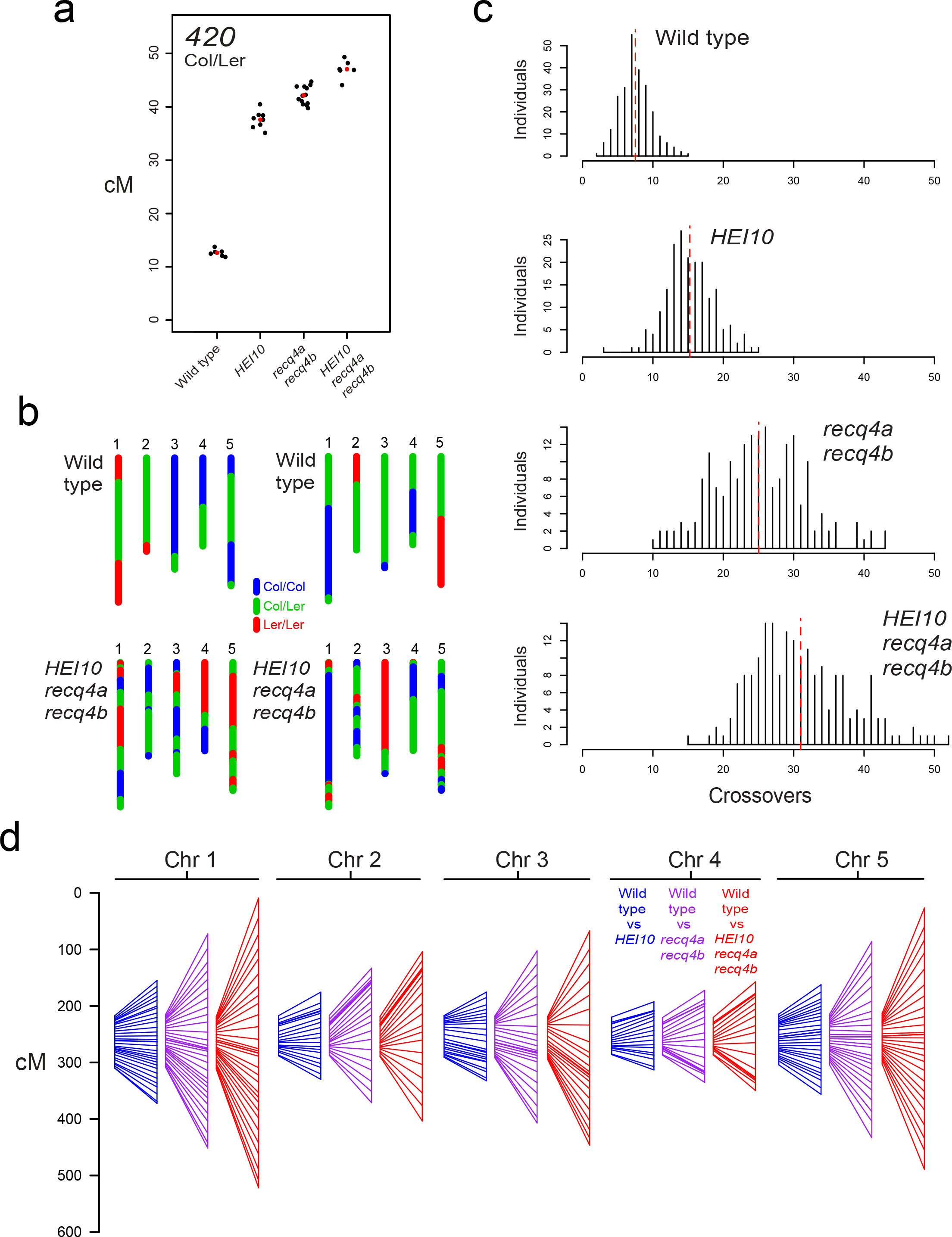
Combination of *HEI10* and *recq4a recq4b* massively increases meiotic crossover frequency. **a.** *420* genetic distance (cM) was measured during breeding of the *HEI10* and *recq4a recq4b* mutant populations. All samples were Col/Ler heterozygous. Replicate measurements are shown as black dots and mean values as red dots. Plotted data is shown in Supplementary Table 1. The *HEI10* data was previously reported^12^. **b.** Chromosomal genotypes are shown from two representative individuals from the wild type and *HEI10 recq4a recq4b* F_2_ populations. The five Arabidopsis chromosomes are depicted and colour-coded according to Col/Col (blue), Col/Ler (green) or Ler/Ler (red) genotypes. **c.** Histograms showing the frequency of F_2_ individuals containing different crossover numbers in each population, with the mean value indicated by the horizontal dotted red lines. **d.** Genetic maps (cM) shown for each chromosome for *HEI10* (blue), *recq4a recq4b* (magenta) and *HEI10 recq4a recq4b* (red). Each map shown alongside the wild type map (left), and markers between the maps are connected.

We sequenced genomic DNA from 191–245 Col/Ler F_2_ progeny derived from wild type, *recq4a recq4b* and *HEI10 recq4a recq4b* F_1_ parents (Fig. 1b and Table 1), and compared to a previously described *HEI10* F_2_ population^12^. We observed that *recq4a recq4b* caused 3.3-fold more crossovers genome-wide (25 crossovers/F_2_, 95% confidence interval +/− 0.93), compared with wild type (7.5 crossovers/F_2_, 95% confidence interval +/− 0.28), which is greater than the 2-fold increase previously seen in *HEI10* (15.3 crossovers/F_2_, 95% confidence interval +/− 0.49)^12^ (Fig. 2b-2d and Table 1). If the *HEI10* and *recq4a recq4b* crossover increases combined in a purely additive manner then we would expect to see the sum of their crossover differentials in *HEI10 recq4a recq4b*, equivalent to 7.5 + 7.7 + 17.4 = 32.6 crossovers/F_2_. Indeed, this was similar to the observed value for *HEI10 recq4a recq4b* of 31 crossovers/F_2_ (95% confidence interval +/− 0.97) (Fig. 2b-2d and Table 1). For all populations, the physically largest chromosomes had the longest genetic maps (Supplementary Fig. 1). A subset of wild type and *HEI10 recq4a recq4b* F_2_ individuals were sequenced to higher depth and crossover patterns found to be robustly identified (Supplementary Fig. 2 and Supplementary Table 2). Together these data show that crossover elevations caused by increased *HEI10* dosage and loss of the *RECQ4A RECQ4B* anti-crossover helicases combine in an additive manner, consistent with class I and class II crossover pathways being independent in Arabidopsis.

### Crossover coincidence increases in both *HEI10* and *recq4a recq4b*

Underdispersion of crossover numbers per meiosis occurs due to the action of crossover interference and homeostasis^6,32^, causing an excess of values close to the mean. Consistently, we observe that the distribution of crossovers per wild type F_2_ individual is significantly non-Poisson (Goodness-of-fit test for Poisson distribution, P=0.012) (Fig. 3a). Observed frequencies are displayed as bars (grey) plotted from the fitted frequencies (red line), such that grey bars lying above or below zero on the y-axis represent deviation from the Poisson expectation (Fig. 3a). Crossover distributions per individual in *recq4a recq4b* and *HEI10 recq4a recq4b* were also non-Poisson (*recq4a recq4b P*=1.85×10^−5^ and *HEI10 recq4a recq4b P=0.0223*). However, the high recombination populations also showed significantly greater variation in crossover numbers compared to wild type (Brown-Levene test, *HEI10 P*=4.17×10^−7^, *recq4a recq4b P*=<2.2× 10^−16^, *HEI10 recq4a recq4b P*=<2.2× 10^−16^) (Fig. 3a and Table 1). We therefore sought to examine the distributions of crossovers within the GBS data in more detail, with respect to inter-crossover spacing.

**Figure 3.**
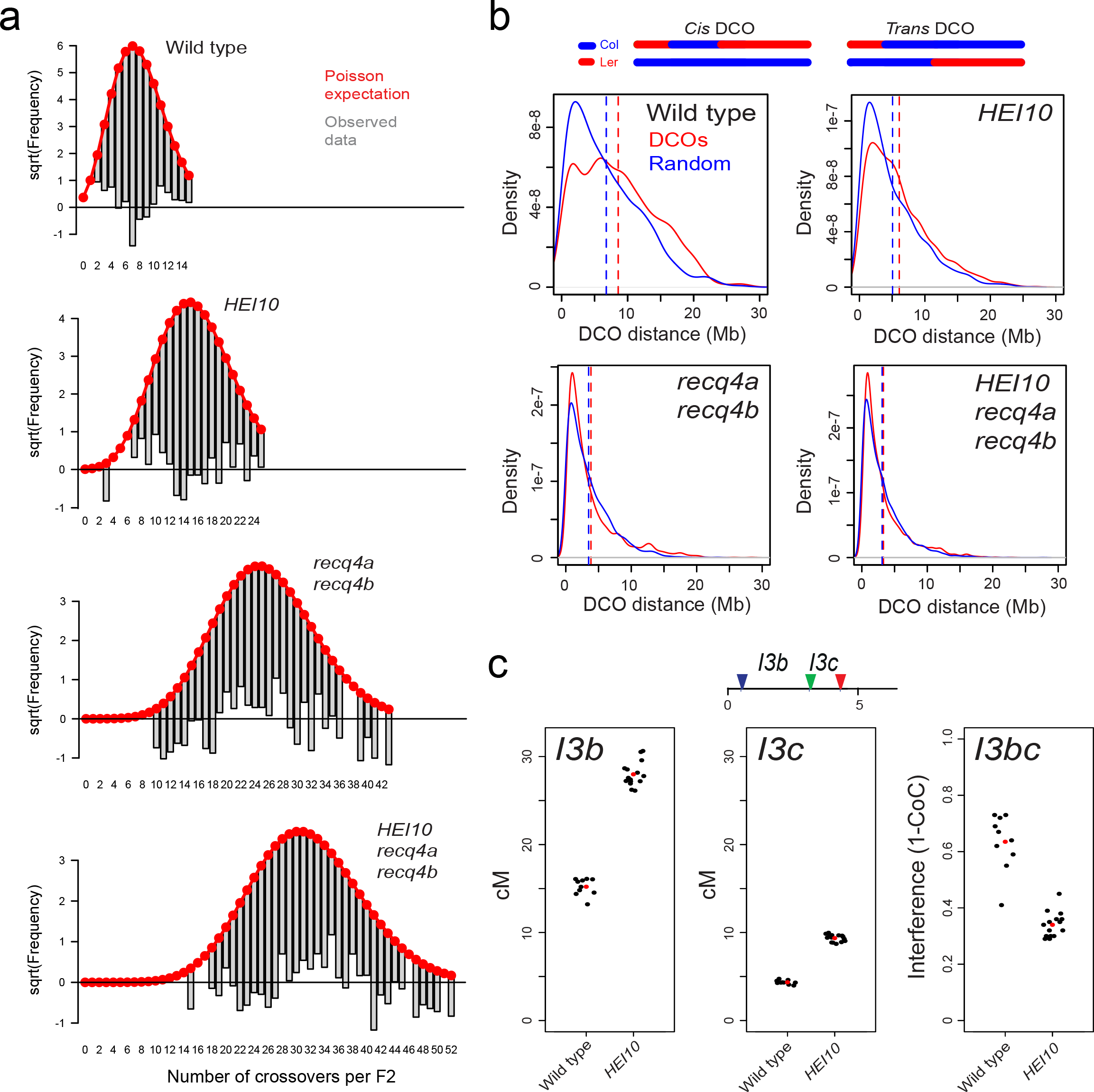
Crossover coincidence increases in *HEI10* and *recq4a recq4b*. **a.** Plots of the square root of the frequency of total crossovers per F_2_ individual in wild type, *HEI10, recq4a recq4b* and *HEI10 recq4a recq4b* populations, generated using the R package goodfit. The expected Poisson distribution is plotted in red, from which the observed data is plotted. Deviation from the Poisson expectation is shown by the grey bay (observed data) falling either above or below the zero value on the y-axis. **b.** A graphical diagram is shown to illustrate *cis* versus *trans* double crossovers detected in F_2_ genotyping-by-sequencing data. Kernel density estimates are plotted for measured DCO distances in the indicated populations (red), versus the same number of matched randomly chosen double events (blue). The vertical dotted lines indicate mean values. **c.** *I3b* and *I3c* genetic distances in wild type and *HEI10*, and crossover interference (1-CoC) between the *I3b* and *I3c* intervals. Replicate measurements are shown by black dots and mean values by red dots. The plotted data in found in Supplementary Tables 3 and 4.

Due to F_2_ individuals being analysed, which result from two independent meioses, it is not possible to distinguish between double crossovers (DCOs) occurring on the same (*cis*) or different (*trans*) chromosomes (Fig. 3b). Importantly, only the *cis* class is informative for estimation of interference. Despite this limitation, we considered each F_2_ individual separately and identified adjacent DCOs for each chromosome and recorded their distances (Fig. 3b). Simultaneously, for each individual and chromosome the same number of randomly chosen positions were used to generate a matched set of randomized distances (Fig. 3b). Consistent with the action of crossover interference, wild type DCO distances were significantly greater than random (mean=8.57 Mb vs 6.99 Mb, Mann-Whitney Wilcoxon test *P*=1.87×10^−16^) (Fig. 3b). In *HEI10* DCO distances were substantially reduced compared to wild type, although they were still significantly greater than random (mean=6.09 Mb vs 5.08 Mb, Mann-Whitney Wilcoxon test *P*=9.46× 10^−13^) (Fig. 3b). However, in both *recq4a recq4b* (mean=3.84 Mb vs 3.49 Mb, Mann Whitney Wilcoxon test P=0.268) and *HEI10 recq4a recq4b* populations (mean=3.30 Mb vs 3.08 Mb, Mann Whitney Wilcoxon test *P*=0.165), observed DCO distances were not significantly different from random (Fig. 3b). This is expected due to increased class II crossovers caused by *recq4a recq4b* being randomly distributed^26^. However, the significant reduction in *HEI10* DCO distances was unexpected, due to this gene acting in the interference-sensitive ZMM pathway^9,12^.

To further investigate crossover interference in *HEI10* we used three-colour FTL analysis, using the adjacent *I3b* and *I3c* intervals, which measure crossover frequency in a sub-telomeric region of chromosome 3^31,33^ (*I3bc* is located within the *420* interval described earlier) (Fig. 2a and 3c). We used flow cytometry to measure inheritance of pollen fluorescence in wild type and *HEI10* and calculate *I3b* and *I3c* genetic distances (Fig. 3c and Supplementary Tables 3–4). Both *I3b* and *I3c* showed a significant increase in crossover frequency in *HEI10*, consistent with our previous *420* measurements (*X^2^* test both P=<2.2×10^−16^) (Figs. 2b, 3c and Supplementary Tables 1 and 3). *I3b* and *I3c* genetic distances were used to estimate the number of DCO pollen expected in the absence of interference, using the formula: Expected DCOs=(I3b cM/100)×(I3c cM/100)×total pollen number. The ratio of ‘observed DCOs’ to ‘expected DCOs’ gives the coefficient of coincidence (CoC), and interference is calculated as 1-CoC, such that zero indicates an absence of interference^31,33^ (Fig. 3c and Supplementary Tables 2–3). Consistent with the reduction in DCO distances seen in the *HEI10* GBS data, *I3bc* interference (1-CoC) significantly decreased from 0.64 in wild type to 0.34 in *HEI10 (X^2^* test *P=* <2.2×10^−16^) (Fig. 3c and Supplementary Tables 3–4). These experiments confirm our GBS observations and reveal that although HEI10 functions in the interfering ZMM pathway, higher *HEI10* dosage causes increased crossover coincidence compared to wild type, although not to the degree observed in *recq4a recq4b*^26^ (Fig. 3b).

### Crossover frequency, interhomolog divergence and DNA methylation landscapes

We next sought to analyse crossover distributions along the chromosomes, and relate these patterns to other aspects of genome organization (Fig. 4). On average 7.5 crossovers were observed per wild type F_2_ individual, 5.6 of which occurred in the chromosome arms and 1.9 in the pericentromeric heterochromatin (Supplementary Fig. 3 and Supplementary Table 5). In *HEI10, recq4a recq4b* and *HEI10 recq4a recq4b*, crossovers in the arms increased 2.3, 4.1 and 5-fold, respectively (5.6 -> 13.1 -> 23 -> 28.2 crossovers), whereas the pericentromeres increases of 1.1, 1.1 and 1.5-fold, respectively (1.9 -> 2.1 -> 2.0 -> 2.7 crossovers), were considerably lower (Supplementary Fig. 3 and Supplementary Table 5). Consistent with previous observations^12,27^, we observed that despite massive crossover increases throughout the chromosome arms, *HEI10, recq4a recq4b* and *HEI10 recq4a recq4b* maintain suppression of recombination within the centromeric regions (Fig. 4a). We also observed that a sub-telomeric region on the long arm of chromosome four showed relative suppression of crossovers, specifically in the *recq4a recq4b* and *HEI10 recq4a recq4b* populations (Fig. 4a). This may reflect a lineage-specific sequence rearrangement, such as an inversion, shared among the *recq4a recq4b* backgrounds.

**Figure 4.**
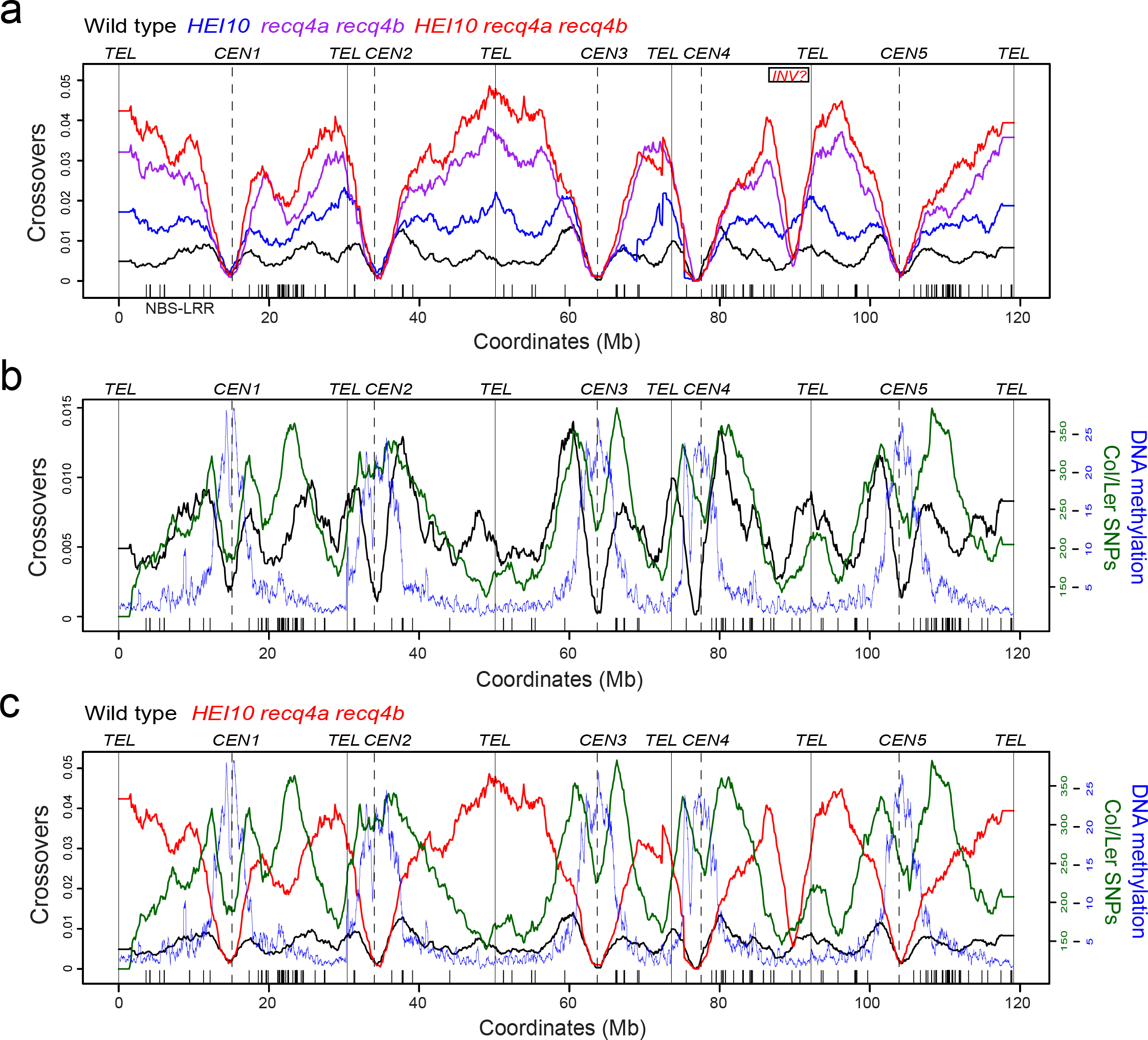
Genomic landscapes of crossover frequency, interhomolog divergence and DNA methylation. **a.**Plots of normalized crossover frequency measured in wild type (black), *HEI10* (blue), *recq4a recq4b* (purple) and *HEI10 recq4a recq4b* (red). The five chromosomes are plotted on a continuous x-axis, with the positions of telomeres (*TEL*) and centromeres (*CEN*) indicated by vertical lines. The position of NBS-LRR resistance gene homologs are indicated by the x-axis ticks. The putative location of an inversion in the *recq4a recq4b* derived populations is also indicated and labelled ‘*INV?*’ **b.** As for a., but showing wild type crossover frequency plotted against Col/Ler SNPs (green)^34^ and DNA methylation (blue)^35^. **c.** As for b., but showing both wild type (black) and *HEI10 recq4a recq4b* (red) crossover frequency.

We hypothesized that genetic and epigenetic factors could contribute to the observed biases in crossover increases towards the telomeres. Therefore, we compared recombination to patterns of Col/Ler interhomolog divergence^34^ (i.e. heterozygosity), and DNA cytosine methylation^35^. Within the chromosome arms we observed that wild type crossovers showed a positive relationship with divergence (r=0.514), which is similar to correlations previously observed between historical recombination and sequence diversity^31^ (Figs. 4b and 5). In contrast, an opposite, negative correlation was seen between *HEI10 recq4a recq4b* crossovers and divergence (r=-0.658) (Figs. 4b and 5). This indicates that the crossover elevations seen in *HEI10 recq4a recq4b* are biased towards the least polymorphic regions of the chromosomes. Hence, while the class II repair pathway that is active in *recq4a recq4b* is not completely inhibited by heterozygosity, it shows a preference for regions of low divergence. The densely DNA methylated centromeric regions are also strongly crossover suppressed in all populations, consistent with heterochromatin inhibiting meiotic recombination^35^ (Figs. 4b and 5). Therefore, although combination of *HEI10* and *recq4a recq4b* causes a massive crossover increase, the localization of recombination is significantly constrained by both interhomolog sequence divergence and chromatin.

**Figure 5.**
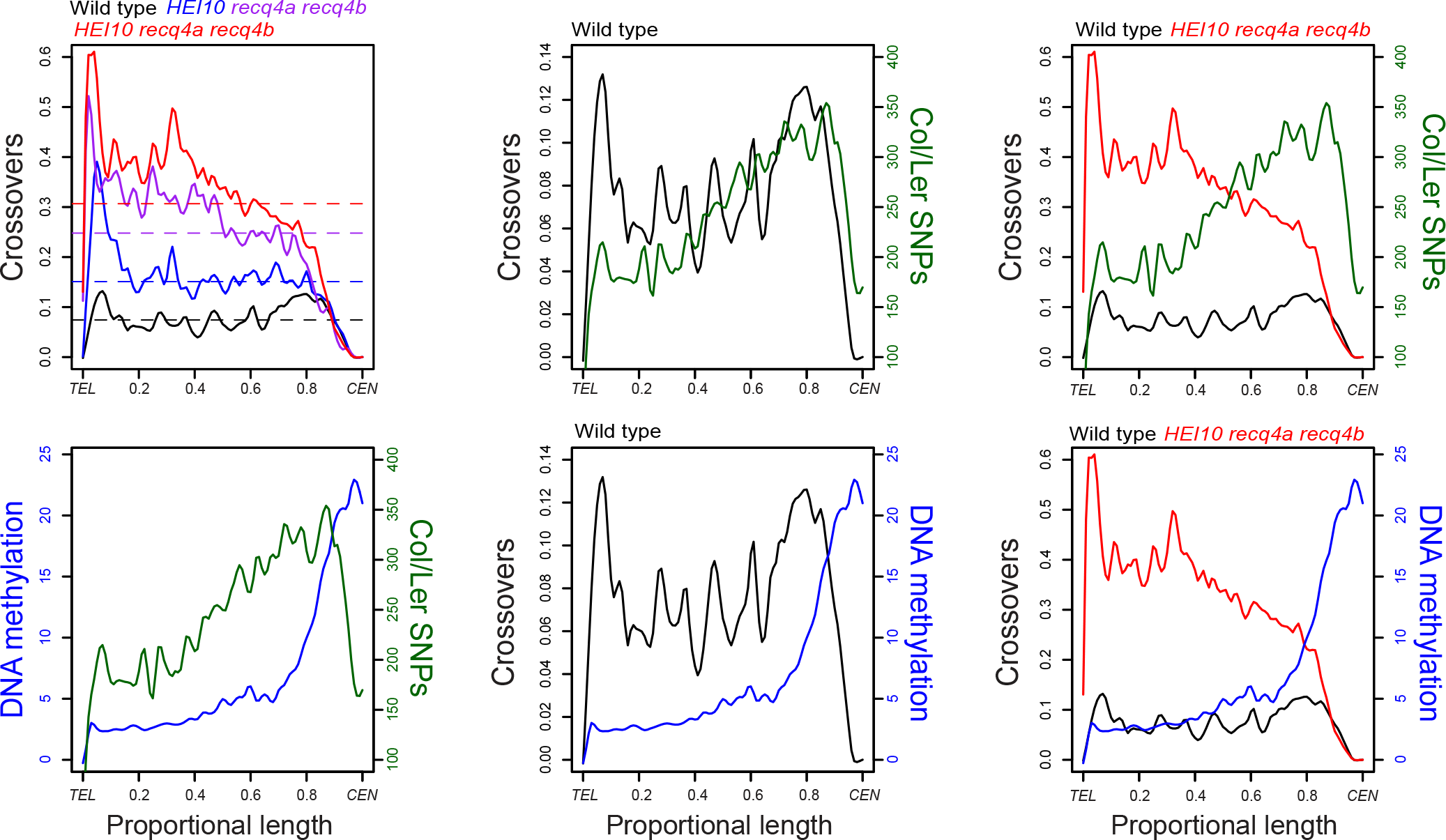
Crossover frequency, interhomolog divergence and DNA methylation along telomere-centromere chromosome axes. Analysis of crossover frequency in wild type (black), *HEI10* (blue), *recq4a recq4b* (purple) and *HEI10 recq4a recq4b* (red), Col/Ler SNPs (green)^34^ and DNA methylation (blue)^35^, analysed along the proportional length of all chromosome arms, orientated from telomeres (*TEL*) to centromeres (*CEN*).

## Discussion

We show that elevating the ZMM crossover pathway, via increased dosage of the *HEI10* meiotic E3 ligase gene, while simultaneously increasing the activity of non-interfering repair, via mutation of *RECQ4A* and *RECQ4B* anti-recombination helicase genes, is sufficient to cause a massive increase in Arabidopsis meiotic crossovers. This is consistent with class I and class II acting as independent crossover repair pathways in plants. HEI10 is a conserved ubiquitin/SUMO E3 ligase with unknown targets during Arabidopsis meiosis, which may include other ZMM factors^1,6,36^. In plants HEI10 associates with paired homologous chromosomes throughout meiotic prophase, showing gradual restriction to a small number of foci that correspond to crossover locations^9,37^. We propose that HEI10 acts quantitatively to promote ZMM pathway crossover repair at recombination sites via SUMO or ubiquitin transfer. Unexpectedly, we show that increased *HEI10* dosage causes higher crossover coincidence and therefore a decrease in genetic interference. Crossover interference has been modelled as a mechanical force, thought to be transmitted via the meiotic chromosome axis and/or synaptonemal complex (SC)^38^. Therefore, HEI10 may modify recombination factors at repair foci and decrease their sensitivity to the interference signal, thereby increasing the likelihood of ZMM-dependent crossover designation. Alternatively HEI10 may alter transmission of the interference signal *per se*, for example, if components of the axis or SC are SUMO/ubiquitin targets.

The RECQ4 helicases have biochemically characterised activities in (i) disassembly of D-loops and (ii) decatenation of dHJs^39–41^, and thus can promote non-crossover outcomes at multiple recombination steps post-strand invasion. In the *recq4a recq4b* mutant it is likely that unrepaired joint molecules persist, which are instead repaired as non-interfering class II crossovers^27^ (Fig. 1a). We show that combination of genetic backgrounds that increase class I and class II crossovers is sufficient to cause a massive and additive recombination increase from 7.5 to 31 crossovers per Arabidopsis F_2_ individual. However, given that ∼100–200 DSB foci have been cytogenetically observed in Arabidopsis there likely remains the capacity for further crossover increases^15–17^. As the Arabidopsis anti-crossover pathways do not show complete redundancy^22,25,27,42^, combination of mutations in the *FANCM*, *RECQ4A-RECQ4B* and *FIGL1* pathways has the potential to cause further increases. Furthermore, the Arabidopsis MSH2 MutS homolog acts to suppress crossovers specifically when homologous chromosomes are polymorphic^43^, and therefore introduction of *msh2* mutations may further increase recombination. The use of *msh2* is attractive, as it may reduce or lessen the bias against crossovers observed in divergent regions in *HEI10 recq4a recq4b*. Equally, modification of epigenetic information has the potential to increase crossovers in centromeric regions. However, as plant heterochromatin is maintained by multiple interacting systems of epigenetic marks, including DNA methylation, H3K9me2, H3K27me1 and H2A.W^44,45^, these marks may have differentiated functions in control of recombination^35^. In conclusion, advanced tailoring of genetic backgrounds may further bias meiotic DSB repair to crossover fates, which has the potential to accelerate crop breeding and improvement.

## Materials and Methods

### Plant Materials

Arabidopsis lines used in this study were the Col *HEI10* line ‘C2’^12^, Col *recq4a-4* (N419423)^29^, Col *recq4b-2* (N511130)^29^ and Ler *recq4a* line (W387*)^27^. Genotyping of *recq4a-4* was performed by PCR amplification using recq4a-F and recq4-wt-R oligonucleotides for wild type and recq4a-F and recq4-mut-R for *recq4a-4*. Genotyping of *recq4b-2* was carried out by PCR amplification using recq4b-wt-F and R oligonucleotides for wild type and recq4b-mut-F and R oligonucleotides for *recq4b-2*. Genotyping of *recq4a* mutation in Ler was performed by PCR amplification using recq4a-Ler-F and R oligonucleotides and subsequent digestion of the PCR products by *Scr*FI restriction enzyme, which yields ∼160 bp products for wild type and ∼180 bp products for *recq4a*. The presence of *HEI10* transgene was tested for by PCR amplification using HEI10-F and HEI10-R oligonucleotides. Oligonucleotide sequences are provided in Supplementary Table 6.

### Measurement of crossover frequency using fluorescent tagged lines

*420* genetic distance was measured using microscopic analysis of seed fluorescence, as described^30,31^. *I3bc* genetic distances and the coefficient of coincidence were measured using fluorescent pollen and flow cytometry, as described^31^.

### Genotyping-by-sequencing

Genomic DNA was extracted and used to generate genotyping-by-sequencing libraries as described^42^. Sequence data was analysed to identify crossover locations using the TIGER bioinformatics pipeline^46^. In order to generate genetic maps the GBS genotypes called from each library were used to call ‘marker’ genotypes at 1 Mb intervals. These calls were then used as an input for the R package Rqtl in order to generate genetic maps using the Haldane mapping function^47^. The R package goodfit was used to compare observed crossover numbers per individual to the Poisson expectation. Statistical analysis of FTL crossover frequency and interference measurements were performed as described^12,31^.

## Acknowledgements

This work was supported by grants from BBSRC (BB/L006847/1), ERA-CAPS/BBSRC ‘DeCOP’ (BB/M004937/1), ERC (CoG), the Gatsby Charitable Foundation (GAT2962) and a Royal Society University Research Fellowship.

